# Repeated duplication of Argonaute2 is associated with strong selection and testis specialization in *Drosophila*

**DOI:** 10.1101/046490

**Authors:** Samuel H. Lewis, Claire L. Webster, Heli Salmela, Darren J. Obbard

**Affiliations:** Institute of Evolutionary Biology, University of Edinburgh, Ashworth Laboratories, EH9 3FL, United Kingdom; Present Address: Department of Genetics, University of Cambridge, Downing Street, Cambridge, CB2 3EH, United Kingdom; Present Address: Life Sciences, University of Sussex, United Kingdom; Department of Biosciences, Centre of Excellence in Biological Interactions, University of Helsinki, Helsinki, Finland; Centre for Immunity, Infection and Evolution, University of Edinburgh, Ashworth Laboratories, EH9 3FL, United Kingdom

**Keywords:** Argonaute, RNAi, *Drosophila*, duplication, testis

## Abstract

Argonaute2 (Ago2) is a rapidly evolving nuclease in the *Drosophila melanogaster* RNA interference (RNAi) pathway that targets viruses and transposable elements in somatic tissues. Here we reconstruct the history of Ago2 duplications across the *Drosophila obscura* group, and use patterns of gene expression to infer new functional specialization. We show that some duplications are old, shared by the entire species group, and that losses may be common, including previously undetected losses in the lineage leading to *D. pseudoobscura*. We find that while the original (syntenic) gene copy has generally retained the ancestral ubiquitous expression pattern, most of the novel Ago2 paralogues have independently specialized to testis-specific expression. Using population genetic analyses, we show that most testis-specific paralogues have significantly lower genetic diversity than the genome-wide average. This suggests recent positive selection in three different species, and model-based analyses provide strong evidence of recent hard selective sweeps in or near four of the six D. pseudoobscura Ago2 paralogues. We speculate that the repeated evolution of testis-specificity in obscura group Ago2 genes, combined with their dynamic turnover and strong signatures of adaptive evolution, may be associated with highly derived roles in the suppression of transposable elements or meiotic drive. Our study highlights the lability of RNAi pathways, even within well-studied groups such as *Drosophila*, and suggests that strong selection may act quickly after duplication in RNAi pathways, potentially giving rise to new and unknown RNAi functions in non-model species.

**Supporting Data:** All new sequences produced in this study have been submitted to Genbank as KX016642-KX016771.

## Introduction

Argonaute genes are found in almost all eukaryotes, where they play a key role in antiviral immune defence, gene regulation and genome stability. They perform this diverse range of functions through their role in RNA interference (RNAi) mechanisms, an ancient system of nucleic acid manipulation in which small RNA (sRNA) molecules guide Argonaute proteins to nucleic acid targets through base complementarity (reviewed in Meister 2013). Gene duplication has occurred throughout the evolution of the Argonaute gene family, with ancient duplication events characteristic of some lineages – such as three duplications early in plant evolution (Singh et al. 2015), and multiple expansions and losses throughout the evolution of nematodes (reviewed in Buck and Blaxter 2013) and the Diptera (Lewis et al. 2016). After duplication, Argonautes have often undergone functional divergence, involving changes in expression patterns and altered sRNA binding partners (Lu et al. 2011; Leebonoi et al. 2015; Miesen et al. 2015). Duplication early in eukaryotic evolution produced two distinct Argonaute subfamilies, Ago and Piwi, which have since been retained in the vast majority of Metazoa (Cerutti and Casas-Mollano 2006). Members of the Ago subfamily are expressed in both somatic and germline tissue, and variously bind sRNAs derived from host transcripts (miRNAs, endo-siRNAs) or transposable elements (TE endo-siRNAs) and viruses (viRNAs). In contrast, in most vertebrates and arthropods, the Piwi subfamily members are expressed primarily in association with the germline (reviewed in Ross et al. 2014), and bind sRNAs from TEs and host loci (piRNAs), suggesting that the Piwi subfamily specialised to a germline-specific role on the lineages leading to vertebrates and arthropods.

After the early divergence of the Ago and Piwi subfamilies, subsequent duplications gave rise to three Piwi subfamily members (Ago3, Aubergine (Aub) and Piwi) and two Ago subfamily members (Ago1 & Ago2) in *Drosophila melanogaster*. All three Piwi subfamily genes are associated with the germline and bind Piwi-interacting RNAs (piRNAs) derived from TEs and other repetitive genomic elements: Ago3 and Aub amplify the piRNA signal through the “Ping-Pong” cycle (reviewed in Luteijn and Ketting 2013), and Piwi suppresses transposition by directing heterochromatin formation (Sienski et al. 2012). These functional differences are associated with contrasting selective regimes, with Aub evolving under positive selection (Kolaczkowski et al. 2011) and more rapidly than Ago3 and Piwi (Obbard, Gordon, et al. 2009). In contrast, Ago1 binds microRNAs (miRNAs), and regulates gene expression by inhibiting translation and marking transcripts for degradation (reviewed in Eulalio et al. 2008). This function imposes strong selective constraint on Ago1, resulting in slow evolution and very few adaptive substitutions (Obbard et al. 2006; Obbard, Gordon, et al. 2009; Kolaczkowski et al. 2011). Finally, Ago2 binds small interfering RNAs (siRNAs) from viruses (viRNAs) and TEs (endo-siRNAs), and functions in gene regulation (Wen et al. 2015), dosage compensation (Menon and Meller 2012), and the ubiquitous suppression of viruses (Li et al. 2002; van Rij et al. 2006) and TEs (Chung et al. 2008; Czech et al. 2008). Ago2 also evolves under strong positive selection, with frequent selective sweeps (Obbard et al. 2006; Obbard, Gordon, et al. 2009; Obbard, Welch, et al. 2009; Kolaczkowski et al. 2011; Obbard et al. 2011), possibly driven by an arms race with virus-encoded suppressors of RNAi (VSRs) (Obbard et al. 2006; Marques and Carthew 2007; van Mierlo et al. 2014).

In contrast to *D. melanogaster*, from which most functional knowledge of Ago2 in 1 arthropods is derived, an expansion of Ago2 has been reported in *D. pseudoobscura* (Hain et al. 2010), providing an opportunity to study how the RNAi pathway evolves after duplication. Given the roles of *D. melanogaster* Ago2 in antiviral defence (Li et al. 2002; van Rij et al. 2006), TE suppression (Chung et al. 2008; Czech et al. 2008), dosage compensation (Menon and Meller 2012), and gene regulation (Wen et al. 2015), we hypothesized that these *D. pseudoobscura* Ago2 paralogues may have diverged in function. To elucidate the evolution and function of Ago2 paralogues in *D. pseudoobscura* and its relatives, we identified and dated Ago2 duplication events across available *Drosophila* genomes and transcriptomes, tested for divergence in expression patterns between the Ago2 paralogues in *D. subobscura*, *D. obscura* and *D. pseudoobscura*, and quantified the evolutionary rate and positive selection acting on each of these paralogues. We find that testis-specificity of Ago2 paralogues has evolved repeatedly in the *obscura* group, and that the majority of paralogues show evidence of recent positive selection.

## Materials and Methods

### Identification of Ago2 homologues in the Drosophilidae

We used tBLASTx to identify Ago2 homologues in transcriptomes and genomes of species of the Drosophilidae, using previously-characterised Ago2 from the closest possible relative to provide the query for each species. If blast returned partial hits, we aligned all hits from the target species to all Argonautes from the query species, and assigned hits to the appropriate Ago lineage based on a neighbour-joining tree. For each query sequence, we then manually curated partial blast hits into complete genes using Geneious v5.6.2 (http://www.geneious.com, Kearse et al. 2012) (see Supplementary Materials for sequence accessions). Additionally, we used degenerate PCR to identify Ago2 paralogues in *D. azteca* and *D. affinis*, and paralogue-specific PCR with a touchdown amplification cycle to validate the Ago2 paralogues identified in *D. subobscura*, *D. obscura* and *D. pseudoobscura*. For each reaction, unincorporated primers were removed with ExonucleaseI (New England Biolabs) and 5’ phosphates were removed with Antarctic Phosphatase (NEB). The PCR products were sequenced by Edinburgh Genomics using BigDye V3 reagents on a capillary sequencer (Applied Biosystems), and Sanger sequence reads were trimmed and assembled using Geneious v.5.6.2 (http://www.geneious.com, Kearse et al. 2012). We also used a combination of PCR and blast searches to locate *D. pseudoobscura* Ago2a1 & Ago2a3, which lie on the unplaced “Unknown_contig_265” in release 3.03 of the *D. pseudoobscura* genome (all PCR primers are detailed in Table S4).

### Phylogenetic analysis of drosophilid Ago2 paralogues

To characterise the evolutionary relationships between Ago2 homologues in the Drosophilidae, we aligned sequences using translational MAFFT (Katoh et al. 2002) with default parameters. We noted that there is a high degree of codon usage bias (CUB) in *D. pseudoobscura* Ago2e (effective number of codons (ENC)=34.24) and *D. obscura* Ago2e (ENC=40.36), and a lesser degree in *D. subobscura* Ago2f (ENC=45.63) and *D. obscura* Ago2f (ENC=48.39), and comparison with genome-wide patterns of codon usage bias placed these genes in the lower half of the distribution of ENC (Figure S5). To reduce the impact of CUB, which disproportionately affects synonymous sites, we stripped all third position sites in this analysis (Behura and Severson 2013). We then inferred a gene tree using the Bayesian approach implemented in BEAST v1.8.1 (Drummond et al. 2012) under a nucleotide model, assuming a GTR substitution model, variation between sites modelled by a gamma distribution with four categories, and base frequencies estimated from the data. We used the default priors for all parameters, except tree shape (for which we specified a birth-death speciation model) and the date of the *Drosophila-Sophophora* split. To estimate a timescale for the tree, we specified a normal distribution for the date of this node using values based on mutation rate estimates in Obbard et al. 2012, with a mean value of 32mya, standard deviation of 7mya, and lower and upper bounds of 15mya and 50mya respectively. We ran the analysis for 50 million steps, recording samples from the posterior every 1,000 steps, and inferred a maximum clade credibility tree with TreeAnnotator v1.8.1 (Drummond et al. 2012). Note that precise date estimates are not a primary focus of this study, but that other calibrations (Russo et al. 1995; Tamura 2004) would lead to more ancient estimates of divergence, and thus stronger evidence for selective maintenance.

### Domain architecture and structural modelling of Ago2 paralogues in the *obscura* group

To infer the location of each domain in each paralogue identified in *D. subobscura*, *D. obscura* and *D. pseudoobscura*, we searched the Pfam database (Finn et al. 2009). To test for structural differences between the *D. pseudoobscura* paralogues, we built structural models of each paralogue based on the published X-ray crystallographic structure of human Ago2 (Schirle and Macrae 2012). We used the MODELER software in the Discovery Studio 4.0 Modeling Environment (Accelrys Software Inc., San Diego, 2013) to calculate ten models, selected the most energetically favourable for each protein, and assessed model quality with the 3D-profile option in the software. To assess variation in selective pressure across the structure of each paralogue, we mapped variable residues onto each structure (Figure S7) using PyMol v.1.7.4.1 (Schrödinger, LLC).

### Quantification of virus-induced expression of Ago2 paralogues

We exposed 48-96hr post-eclosion virgin males and females of *D. melanogaster*, *D. subobscura*, *D. obscura* and *D. pseudoobscura* to *Drosophila C virus* (DCV), by puncturing the thorax with a pin contaminated with DCV at a dose of approximately 4x10^7^ TCID^50^ per ml. Infection with DCV using this method has previously been shown to lead to a rapid and ultimately fatal increase in DCV titre in *D. melanogaster* and *obscura* group species (Longdon et al. 2015). All flies were incubated at 18C under a 12L:12D light cycle, with *D. melanogaster* on Lewis medium and *D. subobscura*, *D. obscura* and *D. pseudoobscura* on banana medium. We sampled 4-7 individuals per species at 0, 8, 16, 24, 48 and 72 hours post infection. At each time-point we extracted RNA using TRIzol reagent (Ambion) and a chloroform/isopropanol extraction, treated twice with TURBO DNase (Ambion), and reverse-transcribed using M-MLV reverse transcriptase (Promega) primed with random hexamers. We then quantified the expression of Ago2 paralogues in these samples by qPCR, using Fast Sybr Green (Applied Biosystems) and custom-designed paralogue-specific qPCR primer pairs (see Table S6 for primer sequences). Due to their high level of sequence similarity (99.9% identity), no primer pair could distinguish between *D. pseudoobscura* Ago2a1 and Ago2a3, so combined expression of these two genes is presented as “Ago2a”. All qPCR reactions for each sample were run in duplicate, and scaled to the internal reference gene Ribosomal Protein L32 (RpL32). To capture the widest possible biological variation, the three biological replicates for each species each used a different wild-type genetic background (see Table S3 for backgrounds used).

### Quantification of Ago2 paralogue expression in different tissues and life stages

For *D. subobscura*, *D. obscura* and *D. pseudoobscura*, we extracted RNA from the head, testis/ovaries and carcass of 48-96hr post-eclosion virgin adults, with males and females extracted separately. Each sample consisted of 8-15 individuals in *D. subobscura*, 10 individuals in *D. obscura* and 15 individuals in *D. pseudoobscura*. We then used qPCR to quantify the expression of each Ago2 paralogue in each tissue, with two technical replicates per sample (reagents, primers and cycling conditions as above). We carried out five replicates per species, each using a different wild-type background (see Table S3 for details of backgrounds used). To provide an informal comparison with the expression pattern of Ago2 before duplication (an “ancestral” expression pattern), we used the BPKM (bases per kilobase of gene model per million mapped bases) values for Ago2 calculated from RNA-seq data from the body (carcass and digestive system), head, ovary and testis of 4 day old *D. melanogaster* adults by Brown et al. 2014, scaling each BPKM value to the value for RpL32 in each tissue. Due to the design of that experiment, the body data are derived from pooled samples of males and females (Brown et al. 2014).

To quantify expression of Ago2 paralogues in *D. pseudoobscura* embryos, we collected eggs within 30 minutes of laying, and used qPCR to measure the expression of each Ago2 paralogue (reagents and primers as above) in two separate wild-type genetic backgrounds (MV8 and MV10). As above, we estimated an ancestral expression pattern of Ago2 before duplication from the BPKM values for Ago2 in 0-2hr old *D. melanogaster* embryos according to Brown et al. 2014, scaled to the BPKM value for RpL32 in embryos. To determine any changes in the expression of other *D. pseudoobscura* Argonautes (Ago1, Ago3, Aub & Piwi) that are associated with Ago2 duplication, we measured their expression in adult tissues and embryos as detailed above, and compared this with the expression of the Argonautes in *D. melanogaster* as measured by Brown et al. 2014.

### Testing for evolutionary rate changes associated with tissue-specificity of Ago2

We used codeml (PAML v4.4, Yang 1997) to fit variants of the M0 model (a single dn/ds ratio, w) to the 65 drosophilid Ago2 homologues shown in Figure 1. All analyses of sequence evolution excluded the highly-repetitive N-terminal glutamine-rich repeat regions, as these regions are effectively unalignable, and are unlikely to conform to simple models of sequence evolution (Palmer and Obbard 2016). In contrast to the tree topology, which was based on 1^st^ and 2^nd^ positions only, the alignment for the codeml analysis included all positions. To compare the evolutionary rates of ubiquitously expressed and testis-specific Ago2 paralogues, we fitted a model specifying one ω for the Ago2 paralogues that were shown to be testis-specific by qPCR (7 homologues), and another ω for the rest of the tree (58 homologues). We also fitted two models to account for rate variation between the *obscura* group Ago2 subclades. The first model specified a separate ω for the Ago2a subclade (17 homologues), the Ago2e subclade (8 homologues), the Ago2f subclade (5 homologues) and the rest of the tree (35 homologues). The second model additionally incorporated an extra ω specified for the *D. pseudoobscura*-*D. persimilis* Ago2a-Ago2b subclade (3 homologues, all of which are testis-specific, in contrast with the rest of the *obscura* group Ago2a subclade). We used Akaike weights to assess which model provided the best fit to the data, given the number of parameters. As mentioned above, the high CUB seen in some Ago2 paralogues may affect PAML analyses by decreasing synonymous site divergence (ds) in those lineages, thereby inflating the dn/ds ratio (w). However, we find no link between levels of codon usage bias and the value of w, suggesting that codon usage bias is not impacting our PAML analyses.

**Figure 1:**
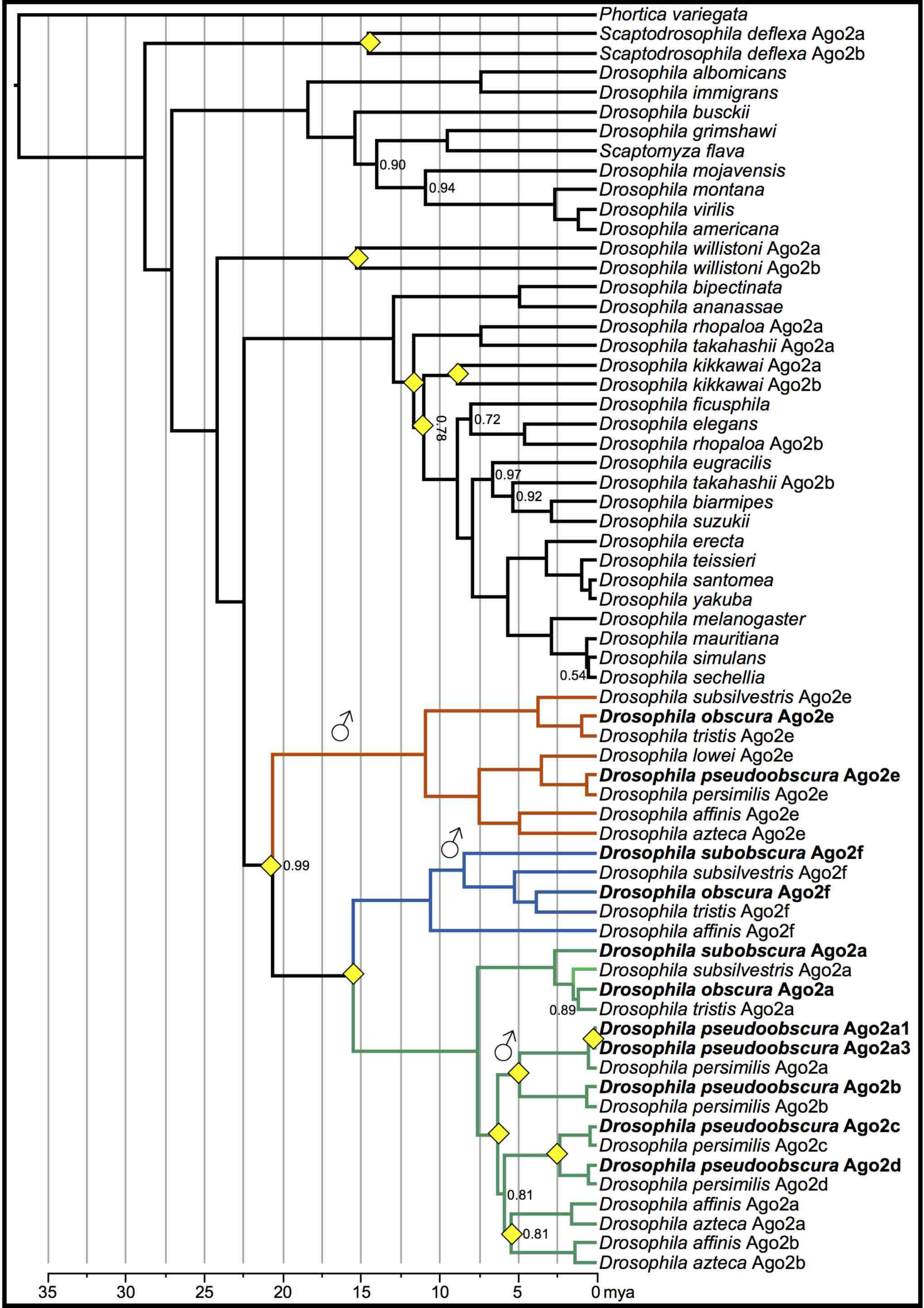
An approximately time-scaled Bayesian gene tree of Ago2 in the Drosophilidae. Duplication events are marked by yellow diamonds, Bayesian posterior support is shown for nodes for which it is less than 100%, and the genes and species that are the focus of the present study are marked in bold. Ago2 has duplicated at least twelve times in the Drosophilidae: seven times in the *obscura* group, twice early in the *melanogaster* group, and once each in the lineages leading to *D. willistoni*, *S. deflexa* and *D. kikkawai*. There has also been a potentially recent duplication of Ago2a on the *D. affinis*/*D. azteca* lineage (∼5mya), although the low support for this node may suggest that these paralogues could also nest within the *D. pseudoobscura*/*D. persimilis* expansion, with one paralogue sister to the Ago2a-Ago2b subclade and the other sister to the Ago2c-Ago2d subclade. After duplication, Ago2 paralogues in the *obscura* group have specialised to the testis three times independently (marked with ♂), and have been retained for an extended period of time (>10 My in the case of Ago2e), suggesting an adaptive basis for testis-specificity. The labelling a-e of paralogous clades corresponds to Hain et al. 2010, and is retained for consistency with subsequent publications which also use these labels, while clade f is newly reported here. All genes were identified by BLAST, apart from the following which were found by PCR: *D. teissieri* Ago2; *D. santomea* Ago2; *D. azteca* Ago2a, Ago2b & Ago2e; *D. pseudoobscura* Ago2a1 & Ago2a3.

### Sequencing of Ago2 paralogue haplotypes from *D. subobscura*, *D. obscura* and *D. pseudoobscura*

To obtain genotype data for the Ago2 paralogues in *D. subobscura*, *D. obscura* and *D. pseudoobscura*, we sequenced the Ago2 paralogues from six males and six females of each species, each from a different wild-collected line (detailed in Table S3, sequence polymorphism data in Appendix S4). We extracted genomic DNA from each individual using the DNeasy Blood and Tissue kit (Qiagen), and amplified and Sanger sequenced each Ago2 paralogue from each individual (reagents and PCR primers as above, sequencing primers detailed in Table S5). We trimmed and assembled Sanger sequence reads using Geneious v.5.6.2 (http://www.geneious.com, Kearse et al. 2012), and identified polymorphic sites by eye. After sequencing Ago2a (annotated as a single gene in the *D. pseudoobscura* genome), we discovered two very recent Ago2a paralogues (which we denote Ago2a1 & Ago2a3), which had been cross-amplified. For each *D. pseudoobscura* individual we therefore re-sequenced Ago2a3 using one primer targeted to its neighbouring locus GA22965, and used this sequence to resolve polymorphic sites in the Ago2a1/Ago2a3 composite sequence, thereby gaining both genotypes for each individual. For each Ago2 paralogue, we inferred haplotypes from these sequence data using PHASE (Stephens et al. 2001), apart from the X-linked paralogues (Ago2a1, Ago2a3 & Ago2d) in *D. pseudoobscura* males, for which phase was obtained directly from the sequence data. The hemizygous haploid X-linked sequenced were used in phase inference, and should substantially improve the inferred phasing of female genotypes.

To quantify differences between paralogues in their population genetic characteristics, we aligned haplotypes using translational MAFFT (Katoh et al. 2002), and used DnaSP v.5.10.01 (Librado and Rozas 2009) to calculate the following summary statistics for each Ago2 paralogue: π (pairwise diversity, with Jukes-Cantor correction as described in Lynch and Crease 1990) at nonsynonymous (π_a_) and synonymous (π_s_) sites, Tajima’s *D* (Tajima 1989) and ENC (Wright 1990). To compare the ENC for each gene with the genome as a whole, we used codonW v1.4.2 (Peden 1995) to calculate the ENC for the longest ORF from each gene or transcript in the genomes or transcriptomes of *D. subobscura*, *D. obscura* and *D. pseudoobscura* (ORF sets detailed below). In each species, we then compared the ENC values of each Ago2 paralogue with this genome-wide ENC distribution.

### Testing for positive selection on Ago2 paralogues in the *obscura* group

We used McDonald-Kreitman (MK) tests (McDonald and Kreitman 1991) to test for positive selection on each Ago2 paralogue. For each paralogue, we chose an outgroup with divergence at synonymous sites (*K_S_*) in the range 0.1-0.2 where possible. However, the prevalence of duplications and losses of Ago2 paralogues in the *obscura* group meant that for some tests no suitably divergent extant outgroup existed. In these cases, we reconstructed hypothetical ancestral sequences using the M0 model provided by codeml from PAML (Yang 1997). To assess the effect of these outgroup choices on our results we repeated each test with another outgroup, and found no effect of outgroup choice on the significance of any tests, and only marginal differences in estimates of α and ω_α_ (results of tests using primary and alternative outgroups are detailed in Table S1 & S2).

A complementary approach to identifying positive selection is to test for reduced diversity at a locus compared with the genome as a whole. To compare the diversity of each *D. pseudoobscura* Ago2 paralogue with the genome-wide distribution of synonymous site diversity, we used genomic data for 12 lines generated by McGaugh et al. 2012. We mapped short reads to the longest ORF for each gene in the R3.2 gene set using Bowtie2 v2.1.0 (Langmead et al. 2009), and estimated synonymous site diversity (θ_W_ based on fourfold synonymous sites) at each ORF using PoPoolation (Kofler et al. 2011). We then plotted the distribution of synonymous site diversity, limited to genes in the size range of 0.75kb - 3kb for comparability with the Ago2 paralogues, and compared the fourfold synonymous site diversity levels of each *D. pseudoobscura* Ago2 paralogue with this distribution. Some *D. pseudoobscura* paralogues are located on autosomes (Ago2b, Ago2c & Ago2e) and some on the X chromosome (Ago2a1, Ago2a3 & Ago2d). Therefore, because of the different population genetic expectations for autosomal and X-linked genes (Vicoso and Charlesworth 2006), we examined separate distributions for autosomal and X-linked genes. To provide an additional test for reduced diversity at *D. pseudoobscura* Ago2 paralogues, we performed maximum-likelihood Hudson-Kreitman-Aguadé tests (Wright and Charlesworth 2004), using divergence from *D. affinis* and intraspecific polymorphism data for 84 *D. pseudoobscura* loci generated by Haddrill et al. 2010. We performed 63 tests to encompass all one, two, three, four, five and six-way combinations of the paralogues, and calculated Akaike weights from the resulting likelihood estimates to provide an estimate of the level of support for each combination.

To infer a genome-wide distribution of synonymous site diversity for *D. obscura* and *D. subobscura*, for which genomic data are unavailable, we used pooled transcriptome data from wild-collected adult male flies that had previously been generated for surveys of RNA viruses (van Mierlo et al. 2014; Webster et al. 2016). To generate a *de novo* transcriptome for each species, we assembled short reads with Trinity r20140717 (Grabherr et al. 2011). For each species, we mapped short reads from the pooled sample to the longest ORF for each transcript, estimated synonymous site diversity at each locus using PoPoolation (Kofler et al. 2011), and plotted the distribution of diversity (as described above for *D. pseudoobscura*). The presence of heterozygous sites in males (identified by Sanger sequencing) confirmed that all Ago2 paralogues in *D. subobscura* and *D. obscura* are autosomal: we therefore compared the synonymous site diversity for these paralogues with the autosomal distribution, and do not show the distributions for putatively X-linked genes. Our use of transcriptome data for *D. obscura* and *D. subobscura* will bias the resulting diversity distributions in three ways. First, variation in expression level will cause individuals displaying high levels of expression to be overrepresented among reads, downwardly biasing diversity. Second, highly expressed genes are easier to assemble, and highly expressed genes tend to display lower genetic diversity (Pal et al. 2001; Lemos et al. 2005). Third, high-diversity genes are harder to assemble, *per se*. However, as all three biases will tend to artefactually reduce diversity in the genome-wide dataset relative to Ago2, this makes our finding that Ago2 paralogues display unusually low diversity conservative.

### Identifying selective sweeps in Ago2 paralogues of *D. pseudoobscura*

To test whether the unusually low diversity seen in the *D. pseudoobscura* Ago2 paralogues is due to recent selection or generally reduced diversity in that region of the genome, we compared diversity at each paralogue to diversity in their neighbouring regions. We obtained sequence data for the 50kb either side of each of these paralogues from the 11 whole genomes detailed in McGaugh et al. 2012 (SRA044960.1, SRA044955.2 & SRA044956.1). Note that the very high similarity of these Ago2 paralogues means that they cannot be accurately assembled from short read data, and are not present in the data from McGaugh et al. 2012. For each genome, we therefore replaced the poorly-assembled region corresponding to the paralogue with one of our own Sanger-sequenced haplotypes, making a set of 11 ca. 102kb sequences for each paralogue. We aligned these sequences using PRANK (Löytynoja and Goldman 2005) with default settings, and calculated Watterson’s θ at all sites in a sliding window across each alignment, with a window size of 5kb and a step of 1kb. For Ago2a1 and Ago2a3, which are located in tandem, we analysed the same genomic region. Since our Ago2 haplotypes were sampled from a different North American population of *D. pseudoobscura* to those of McGaugh et al. 2012, an apparent reduction in local diversity might result from differences in diversity between the two populations. We therefore also repeated these analyses on a dataset in which our Sanger sequenced haplotypes were removed, leaving missing data.

To test explicitly for selective sweeps at each region, we used Sweepfinder (Nielsen, Williamson, et al. 2005) to calculate the likelihood and location of a sweep in or near each Ago2 paralogue. We specified a grid size of 20,000, a folded frequency spectrum for all sites, and included invariant sites. To infer the significance of any observed peaks in the composite likelihood ratio, we used ms (Hudson 2002) to generate 1000 samples of 11 sequences under a neutral coalescent model. We generated separate samples for each region surrounding an Ago2 paralogue, conditioning on the number of polymorphic sites observed in that region, the sequence length equal to the alignment length, and an effective population size of 10^6^ (based on a previous estimate for *D. melanogaster* by Li and Stephan 2006). We specified the recombination rate at 5cM/Mb, a conservative value based on previous estimates for *D. pseudoobscura* (McGaugh et al. 2012), which will lead to larger segregating linkage groups and therefore a more stringent significance threshold.

## Results

### Ago2 has undergone numerous ancient and recent duplications in the *obscura* group

Ago2 duplications had previously been noted in *D. pseudoobscura* (Hain et al. 2010), but their age and distribution in other species was unknown. We used BLAST (Altschul et al. 1997) and PCR to identify 65 Ago2 homologues in 39 species sampled across the Drosophilidae, including 30 homologues in 9 *obscura* group species. Using PCR and Sanger sequencing, we verified that the paralogues in *D. subobscura*, *D. obscura* and *D. pseudoobscura* are genuine distinct loci, and not artefacts of erroneous assembly. Additionally, we verified that all paralogues possess introns, and so are most likely to be the product of segmental duplication rather than retrotransposition. This is perhaps unsurprising given that segmental duplicates are generally retained at a higher rate than retrotransposed duplicates, despite the rate of retrotransposition being higher than segmental duplication (Hahn, 2009).

To characterize the relationships between Ago2 homologues in the *obscura* group and the other Drosophilidae, and estimate the date of the duplication events that produced them, we carried out a strict clock Bayesian phylogenetic analysis (Figure 2). This showed that there are early diverging Ago2 clades in the *obscura* group: the Ago2e subclade that diverged from other Ago2 paralogues around 21mya (±10 My), and the Ago2a and Ago2f subclades that were produced by a gene duplication event around 16mya (±7 My). Subsequently there have been a series of more recent duplications in the *D. pseudoobscura* subgroup Ago2a-d lineage. Using published genomes, transcriptomes and PCR we were unable to identify Ago2e in *D. subobscura*, Ago2e or Ago2f in *D. lowei*, or Ago2f in *D. pseudoobscura*, *D. persimilis* and *D. azteca*. While apparent losses may reflect a lack of genomic data (*D. subobscura*, *D. lowei* and *D. azteca*), incomplete genome assemblies ( *D. pseudoobscura* and *D. persimilis*) or unexpressed genes in transcriptome surveys, we attempted to validate the losses observable in *D. pseudoobscura* and *D. subobscura* by extensive PCR, and were again unable to recover these genes from those two species.

**Figure 2:**
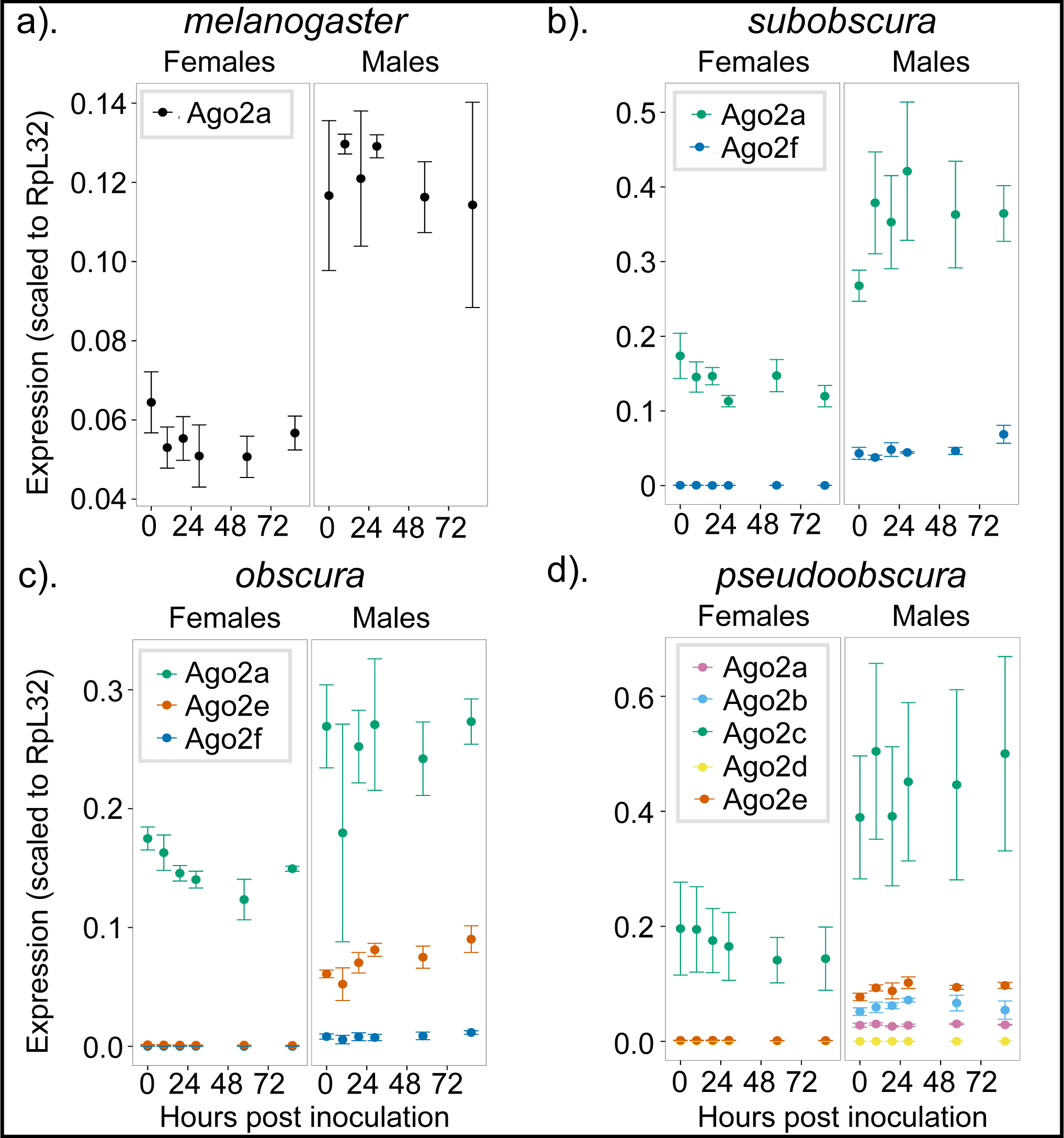
Expression patterns of Ago2 paralogues under challenge with *Drosophila C Virus*. In each *obscura* group species, only one Ago2 paralogue has retained the ancestral pattern of ubiquitous stable expression in each sex (illustrated by *D. melanogaster*). In contrast, all other paralogues are expressed in males only (in *D. pseudoobscura* females, Ago2a, Ago2b, Ago2d & Ago2e are all unexpressed throughout the timecourse). The only exception to this is *D. pseudoobscura* Ago2d, which is unexpressed in either sex. The high degree of sequence similarity between Ago2a1 and Ago2a3 prevented us from amplifying these genes separately in qPCR, and here they are combined as “Ago2a”. Error bars indicate standard error estimated from technical replicates in each of three different genetic backgrounds. Apparent differences in expression between sexes and species should be interpreted with caution, as these may be driven by differences in expression levels of the reference gene (RpL32).

In release 3.03 of the *D. pseudoobscura* genome the paralogues Ago2b-Ago2e have confirmed locations, but Ago2a1 and Ago2a3 (the very recent paralogues newly identified here) lie in tandem on an unplaced contig with a third incomplete copy (Ago2a2) between them. We used PCR to confirm the existence, orientation, and relative positioning of these genes, and to identify the location of this contig, which lies in reverse orientation on chromosome XL-group1a (predicted coordinates 3,463,701-3,489,689). We then combined this information with our phylogenetic analysis to reconstruct the positional evolution of *D. pseudoobscura* Ago2 paralogues (Figure S1). We found that *D. pseudoobscura* Ago2d is syntenic with *D. melanogaster* Ago2, indicating that Ago2d is the ancestral paralogue in this species. We also found that Ago2 paralogues have translocated throughout the *D. pseudoobscura* genome (Figure S1), and are situated on autosomes (Ago2b, Ago2c & Ago2e) and both arms of the X chromosome (Ago2a1, Ago2a3 & Ago2d). It should be noted that a lack of genomic data precludes similar synteny analysis for any other *obscura* group species; our naming of the Ago2 paralogues in these species as Ago2a (or Ago2a and Ago2b in the case of *D. affinis* and *D. azteca*) reflects their position within the Ago2a subclade, rather than a syntenic relationship or otherwise with *D. pseudoobscura* Ago2a1 and Ago2a3.

### Ago2 paralogues in *D. subobscura*, *D. obscura* and *D. pseudoobscura* are probably functional

Our phylogenetic analysis (Figure 1) revealed that the Ago2 paralogues in the *obscura* group have retained coding sequences for millions of generations, showing that they have remained functional for this period. They have also retained PAZ and PIWI domains and a bilobal structure (characteristic of Argonaute proteins), suggesting that they are part of a functional RNAi pathway. In *D. melanogaster* Ago2 plays a key role in antiviral immunity, but is ubiquitously and highly expressed in both males and females, and is not strongly induced by viral challenge (Figure 2a, Aliyari et al. 2008). To test whether this expression pattern has been conserved after Ago2 duplication, or whether any Ago2 paralogues have become inducible by viral challenge, we measured the expression of each Ago2 paralogue in female and male *D. subobscura*, *D. obscura* and *D. pseudoobscura* after infection with *Drosophila C Virus* (DCV). These species are separated by ∼10My of evolution, and represent the three major clades within the *obscura* group. Members of the *obscura* group are highly susceptible to DCV, supporting high viral titres and displaying rapid mortality (Longdon et al. 2015). We found that only one paralogue is expressed in both sexes at a high level in *D. subobscura* (Ago2a), *D. obscura* (Ago2a) and *D. pseudoobscura* (Ago2c). These paralogues show a similar pattern of expression to *D. melanogaster* Ago2, being expressed constitutively throughout the timecourse rather than induced by viral infection (Figure 2). Unexpectedly, and with only one exception, the other Ago2 paralogues in all species were expressed exclusively in males (Figure 2b-d), raising the possibility that these duplicates have specialised to a sex-specific role. The one exception was *D. pseudoobscura* Ago2d, which is the ancestral paralogue in this species (inferred by synteny), and for which we could not detect any expression.

### Ago2 paralogues have repeatedly specialised to the testis

To determine whether the strongly male-biased expression pattern is associated with a testis-specific role, we quantified the tissue-specific expression patterns of Ago2 paralogues in *D. subobscura*, *D. obscura* and *D. pseudoobscura*. In *D. melanogaster* the single copy of Ago2 was expressed in all adult tissues (Figure 3d), and transcripts were present in the embryo (Figure S2). In *D. subobscura*, *D. obscura* and *D. pseudoobscura*, we found that the Ago2 paralogues exhibited striking differences in their tissue-specific patterns of expression (Figure 3a-c). In each species, one paralogue has retained the ancestral ubiquitous expression pattern in adult tissues. In contrast, every other paralogue was expressed only in the testis, except for the non-expressed *D. pseudoobscura* Ago2d. None of the testis-specific paralogues in *D. pseudoobscura* was detectable in embryos (Figure S2).

**Figure 3:**
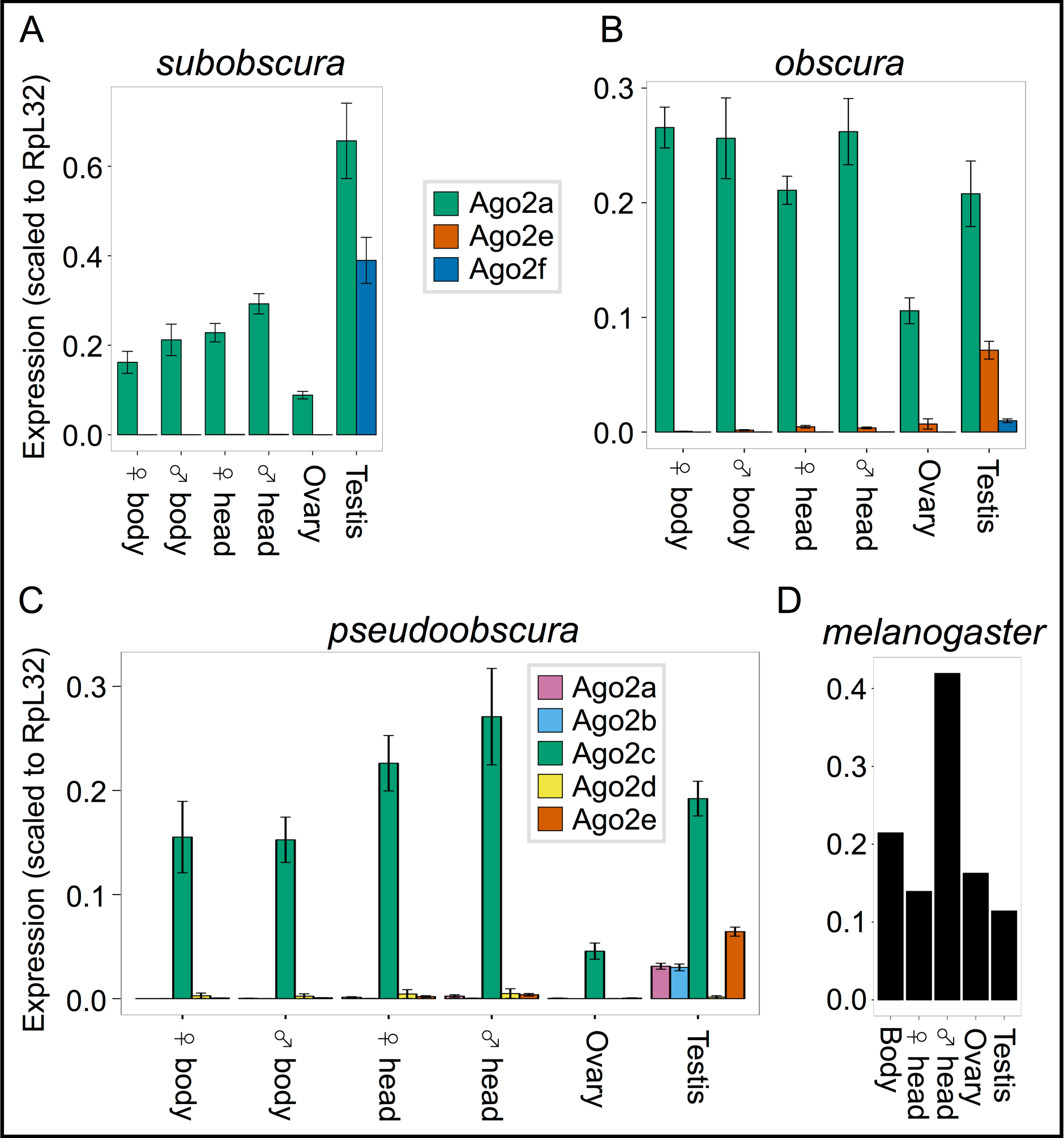
Tissue-specific expression patterns of Ago2 paralogues. In each of the three *obscura* group species tested, one paralogue has retained the ancestral ubiquitous expression pattern, while the others have specialised to the testis (with the exception of *D. pseudoobscura* Ago2d). The high degree of sequence similarity between Ago2a1 and Ago2a3 prevented us from amplifying these genes separately in qPCR, and here they are combined as “Ago2a”. Error bars indicate standard error estimated from technical replicates in each of five different genetic backgrounds. *D. melanogaster* expression levels were taken from a single RNA-seq experiment (Brown et al. 2014).

Interestingly, the ubiquitously expressed paralogue in *D. subobscura* and *D. obscura* is the ancestral gene (Ago2a in both cases, as inferred by synteny with *D. melanogaster*), but in *D. pseudoobscura* another paralogue (Ago2c) has evolved the ubiquitous expression pattern, and the ancestral gene (Ago2d) was not expressed at a detectable level in any tissue. When interpreted in the context of the phylogenetic relationships between these paralogues, the most parsimonious explanation is that testis-specificity evolved at least three times: first at the base of the Ago2e clade, second at the base of the Ago2f clade, and third at the base of the *D. pseudoobscura-D. persimilis* Ago2a-Ago2b subclade (Figure 1).

### Testis-specificity is associated with faster protein evolution

To test for differences in evolutionary rate between testis-specific and ubiquitously expressed Ago2 paralogues, we fitted sequence evolution models to the set of drosophilid Ago2 sequences depicted in Figure 1 using codeml (PAML, Yang 1997). These tests estimate separate dN/dS ratios (w) for different subclades in the gene tree, providing a test for differential rates of protein evolution. We found that most support (Akaike weight = 0.99) falls behind a model specifying a different ω for each obscura group Ago2 subclade, and another separate ω for the *D. pseudoobscura-D. persimilis* Ago2a-Ago2b subclade. Under this model, the testis-specific *D. pseudoobscura*-*D. persimilis* Ago2a-Ago2b subclade has the highest rate of protein evolution (ω=0.32±0.047 SE), followed by the testis-specific Ago2f subclade (ω=0.21±0.014), the ubiquitous Ago2a subclade (ω=0.19±0.012), the testis-specific Ago2e subclade (ω=0.16±0.010), and finally the other Drosophilid Ago2 sequences (ω=0.12±0.002). This shows that the evolution of testis-specificity was accompanied by an increase in the rate of protein evolution following two of the three duplications. We also used the Bayes Empirical Bayes sites test in codeml to identify codons evolving under positive selection across the entire gene tree, and the branch-sites test to identify codons under positive selection in the *obscura* group Ago2 subclade. While we found no positively-selected codons with the sites test, we identified three codons under positive selection (297, 338 & 360) in the *obscura* group Ago2 subclade with the branch-sites test (likelihood ratio test M8 vs M8a, p<0.005).

### McDonald-Kreitman tests identify strong positive selection on *D. pseudoobscura* Ago2e

Changes in evolutionary rate after the evolution of testis-specificity may occur as a result of changes in positive selection, or changes in selective constraint. However, unless there are multiple substitutions within single codons, this will be hard to detect using methods such as codeml. Therefore, as a second test for positive selection on Ago2 paralogues in *D. subobscura*, *D. obscura* and *D. pseudoobscura*, we gathered intraspecies polymorphism data for each Ago2 paralogue in these species (Appendix S4), and performed McDonald-Kreitman (MK) tests (Table S1). The MK test uses a comparison of the numbers of fixed differences between species at nonsynonymous (Dn) and synonymous (Ds) sites, and polymorphisms within a species at nonsynonymous (Pn) and synonymous (Ps) sites to infer the action of positive selection. If all mutations are either neutral or strongly deleterious, the Dn/Ds ratio should be approximately equal to the Pn/Ps ratio; however, if there is positive selection, an excess of nonsynonymous differences is expected (McDonald and Kreitman 1991). The majority of MK tests were non-significant (Fisher’s exact test, p>0.1), despite often displaying relatively high K_A_/K_S_ ratios e.g. *D. pseudoobscura* Ago2a1 (K_A_/K_S_ =0.34), Ago2b (K_A_/K_S_ =0.43) & Ago2d (K_A_/K_S_ =0.36). However, the low diversity at these loci (<10 polymorphic sites in most cases; see below) means that the MK approach has little power, and that estimates of the proportion of substitutions that are adaptive (a) are likely to be poor. In contrast to the other loci, our MK analysis identified strong positive selection acting on *D. pseudoobscura* Ago2e – which has relatively high genetic diversity – with a at 100% (a=1.00; Fisher’s exact test, p=0.0004). This result is driven by the extreme dearth of nonsynonymous to synonymous polymorphisms (0 Pn to 17 Ps), despite substantial numbers of fixed differences (77 Dn to 120 Ds), and its statistical significance is robust to the choice of outgroup (Table S2).

### The majority of Ago2 paralogues have extremely low levels of sequence diversity

When strong selection acts to reduce genetic diversity at a locus, it can also reduce diversity at linked loci before recombination can break up linkage (Maynard Smith and Haigh 1974). Recent positive selection can therefore be inferred from a reduction in synonymous site diversity compared with other genes. Because MK tests can only detect multiple long-term substitutions, and are hampered by low diversity, diversity-based approaches offer a complementary way to detect very recent strong selection. We therefore compared the synonymous site diversity at each Ago2 paralogue in *D. pseudoobscura* with the distribution of genome-wide synonymous site diversity. We found that all *D. pseudoobscura* paralogues have unusually low diversity relative to other loci: Ago2a1, Ago2b and Ago2c fall into the lowest percentile, Ago2a3 and Ago2d into the 2nd lowest percentile and Ago2e into the 8th lowest percentile (Figure S4). A multi-locus extension of the HKA test (ML-HKA, Wright and Charlesworth 2004) confirmed that the diversity of Ago2a1-Ago2e is significantly lower than the *D. pseudoobscura* genome as a whole (Akaike weight = 0.98).

Unfortunately, population-genomic data are not available for *D. subobscura* and *D. obscura*, preventing a similar analysis. However, we found similar results for Ago2a and Ago2e when comparing the diversity of *D. subobscura* and *D. obscura* Ago2 paralogues to levels of diversity inferred from transcriptome data (data from Webster et al. 2016), suggesting that this effect is not limited to *D. pseudoobscura* and these genes may therefore have been recent targets of selection in multiple species. In *D. obscura*, Ago2a and Ago2e fall into the 2nd and 4th lowest diversity percentile respectively, whereas Ago2f falls into the 19th percentile (Figure S4). In *D. subobscura*, Ago2a falls into the 7th percentile, whereas Ago2f falls into the 16th percentile (Figure S4). The prevalence of low intraspecific diversity for testis-specific paralogues is consistent with recent selective sweeps, suggesting that positive selection, not merely relaxation of constraint, has contributed to the increased evolutionary rate seen after specialization to the testis.

### Four out of six *D. pseudoobscura* Ago2 duplicates show a strong signature of recent hard selective sweeps

The impact of selection on linked diversity (a selective sweep) is expected to leave a characteristic footprint in local genetic diversity around the site of selection, and this forms the basis of explicit model-based approaches to detect the recent action of positive selection (Nielsen, Bustamante, et al. 2005). For *D. pseudoobscura*, population genomic data for 11 haplotypes is available from McGaugh et al. 2012, permitting an explicit model-based test for recent hard selective sweeps near to Ago2 paralogues. We therefore combined our Ago2 data with 111kb long haplotypes from McGaugh et al. 2012 to analyse the neighbouring region around each paralogue. Ago2a1 and Ago2a3 form a tandem repeat, and were therefore analysed together as a single potential sweep. We found strong evidence for recent selective sweeps at or very close to Ago2a1/3, Ago2b and Ago2c, which display sharp troughs in their diversity levels, and large peaks in the composite likelihood of a sweep, which far exceed a significance threshold derived from coalescent simulation (p<0.01; Figure 4). These localised reductions in diversity remain when our own Ago2 haplotype data are removed, showing the results are robust to the fact that our Ago2 sequence data are derived from a different population to the genome-wide data of McGaugh et al. 2012 (Figure S6; note that sequence data for Ago2 paralogues cannot be derived from the data of McGaugh et al. 2012, because of their extreme similarity). In addition, there is ambiguous evidence for a sweep at Ago2d, in the form of one significant (p<0.01) likelihood peak just upstream of the paralogue, but two other peaks ∼1kb and ∼3kb further upstream. There is no evidence for a hard sweep at Ago2e, which has no diversity trough or likelihood peak.

**Figure 4:**
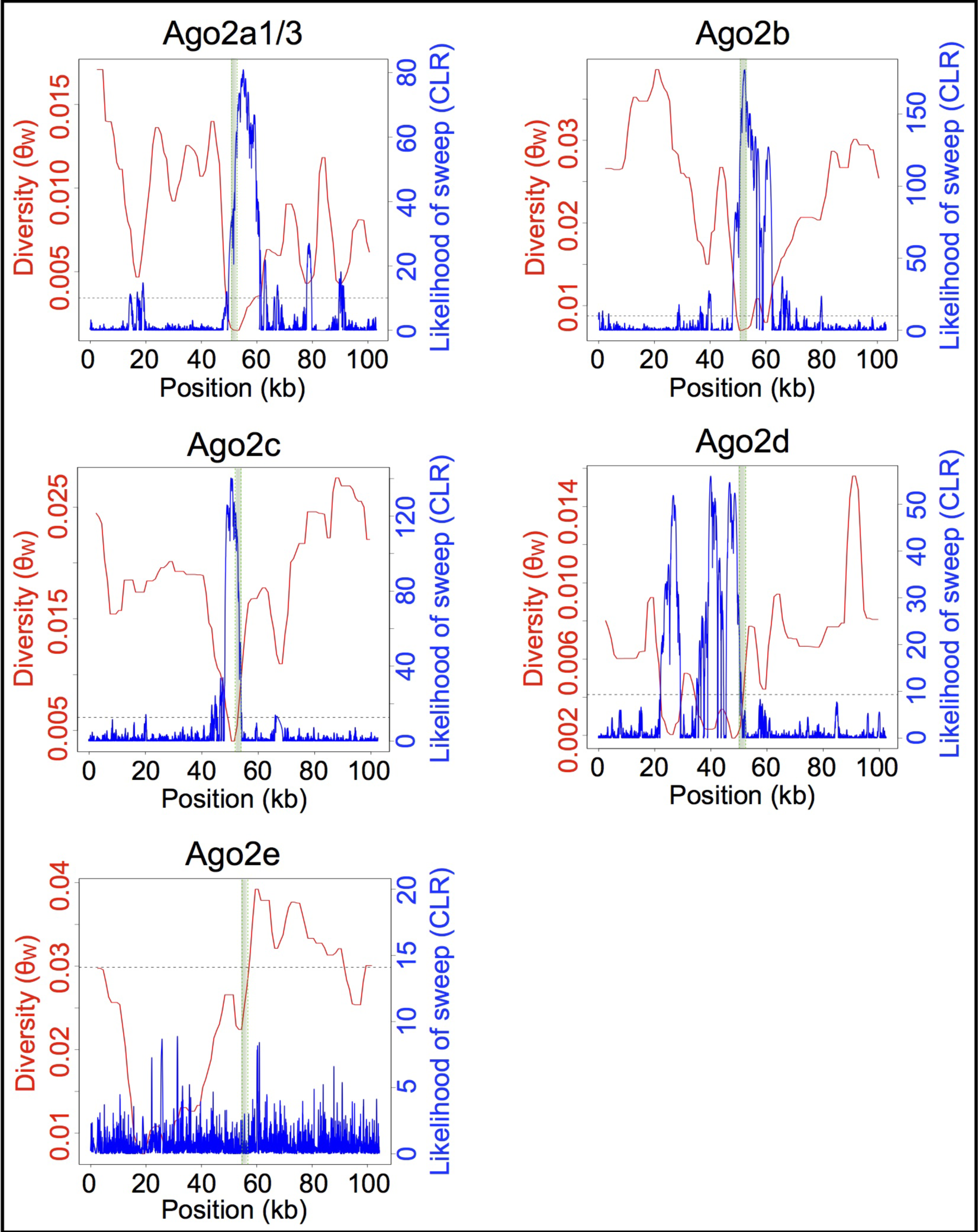
Selective sweeps at *D. pseudoobscura* Ago2 paralogues. For each paralogue, diversity at all sites (Watterson’s θ) is displayed in red, and the likelihood of a sweep centred at that site (composite likelihood ratio, CLR) is displayed in blue. The gene region containing the paralogue is represented by the shaded vertical bar, and the significance threshold for the CLR is displayed by the horizontal dotted line (p<0.01, derived from the 10th-highest CLR out of 1000 coalescent simulations, assuming constant recombination rate and N_e_). There is strong evidence for sweeps at Ago2a, Ago2b and Ago2c, indicated by troughs in their diversity levels and peaks in the likelihood of a sweep.

## Discussion

### Testis-specificity may indicate a loss of antiviral function

We have found that Ago2 paralogues in the *obscura* group have repeatedly evolved divergent expression patterns after duplication, with the majority of paralogues specializing to the testis. This is the first report of testis-specificity for any arthropod Ago2, which is ubiquitously expressed in *D. melanogaster* (Celniker et al. 2009), and provides a strong indication that these paralogues have diverged in function. This testis-specificity (Figure 3) suggests that these Argonautes are likely to have lost their ancestral ubiquitous antiviral role. Additionally, the constant level of expression of testis-specific paralogues under DCV infection (Figure 2) suggests that have not evolved an inducible response to viral infection, either restricted to the testis or in other tissues. In contrast, one paralogue in each species has retained the ubiquitous expression pattern seen in *D. melanogaster* ( *D. subobscura* Ago2a, *D. obscura* Ago2a & *D. pseudoobscura* Ago2c, Figure 3), suggesting that these paralogues have retained roles in antiviral defence (Li et al. 2002; van Rij et al. 2006), dosage compensation (Menon and Meller 2012) and/or somatic TE suppression (Chung et al. 2008; Czech et al. 2008).

### Both ubiquitous and testis-specific Ago2 paralogues show evidence of recent positive selection

We identified selective sweeps at the ubiquitously expressed Ago2 paralogue in *D. pseudoobscura* Ago2c, and very low diversity in the ubiquitously expressed Ago2 paralogues of *D. subobscura* and *D. obscura* (Ago2a), suggesting that all of these genes may have recently experienced strong positive selection. Four randomly-chosen testis-specific genes in *D. obscura* and *D. subobscura* do not fall into the low-diversity tails of the genome-wide diversity distributions, suggesting that this is not a general phenomenon of testis-specific expression. This is consistent with previous findings of strong selection and rapid evolution of Ago2 in *D. melanogaster* (Obbard et al. 2006; Obbard, Welch, et al. 2009; Obbard et al. 2011) which has also experienced recent sweeps in *D. melanogaster*, *D. simulans*, and *D. yakuba* (Obbard et al. 2011), and across the *Drosophila* more broadly (Kolaczkowski et al. 2011). It has previously been suggested that this is driven by arms-race coevolution with viruses (Obbard, Gordon, et al. 2009; Kolaczkowski et al. 2011), some of which encode viral suppressors of RNAi (VSRs) that block Ago2 function (Bronkhorst and van Rij 2014). The presence of VSR-encoding viruses, such as Nora virus, in natural *obscura* group populations (Webster et al. 2016), combined with the host-specificity that can be displayed by VSRs (van Mierlo et al. 2014), suggest that arms-race dynamics may also be driving the rapid evolution of ubiquitously expressed Ago2 paralogues in the *obscura* group.

### Potential testis-specific functions

In contrast to their ancestral ubiquitous expression pattern, the dominant fate for Ago2 paralogues in the *obscura* group appears to have been specialization to the testis. Paralogues often undergo a brief period of testis-specificity soon after duplication (Assis and Bachtrog 2013; Assis and Bachtrog 2015), and this has given rise to the ‘out-of-the-testis’ hypothesis, in which new paralogues are initially testis-specific before evolving functions in other tissues (Kaessmann 2010). However, two lines of evidence suggest an adaptive basis for the testis-specificity observed for the *obscura* group Ago2 paralogues. First, testis-specificity has been retained for more than 10 million years in Ago2e and Ago2f, in contrast to the broadening of expression over time expected under the out-of-the-testis hypothesis (Kaessmann 2010; Assis and Bachtrog 2013). Second, all testis-specific Ago2 paralogues in *D. pseudoobscura* show evidence either of long-term positive selection (MK test for the high-diversity Ago2e) or of recent selective sweeps (in low-diversity Ago2a1/3 and Ago2b), and the testis-specific *D. obscura* Ago2e displays a reduction in diversity, potentially driven by selection.

Under a subfunctionalization model for Ago2 testis-specialization, five candidate selective pressures seem likely: testis-specific dosage compensation, antiviral defence, gene regulation, TE suppression, and/or the suppression of meiotic drive. Of these, testis-specific dosage compensation seems the least likely to drive testis-specificity because the male-specific lethal (MSL) complex, which Ago2 directs to X-linked genes to carry out dosage compensation in the soma of *D. melanogaster*, is absent from testis (Conrad and Akhtar 2012). Testis-specific antiviral defence seems similarly unlikely, as the only known paternally-transmitted *Drosophila* viruses (Sigmaviruses; Rhabdoviridae) pass through both the male and female gametes (Longdon and Jiggins 2012), and so the potential benefits of testis-specificity seem unclear. Alternatively, testis-specific Ago2 duplicates could be co-evolving with other testis-specific genes through the hairpin RNA pathway, in which siRNAs generated from endogenous hairpin-forming RNAs (hpRNAs) bind Ago2 and regulate the expression of host genes (Okamura et al, 2008). In *D. melanogaster*, hpRNA-derived siRNAs target testis-specific genes involved in male fertility, and coevolve with these targets to maintain base complementarity (Wen et al, 2015). If a similar pathway operates in the *obscura* group, Ago2 paralogues could have specialized to the hpRNA pathway in order to regulate testis-specific genes more effectively. Finally, the suppression of TEs or meiotic drive seem promising candidate selective forces. First, numerous TEs transpose preferentially in the testis, such as *Penelope* in *D. virilis* (Rozhkov et al. 2010) and *copia* in *D. melanogaster* (Pasyukova et al. 1997; Morozova et al. 2009), which could impose a selection pressure on Ago2 paralogues to provide a testis-specific TE suppression mechanism. It should be noted that all members of the canonical anti-TE Piwi subfamily (Ago3, Aub and Piwi) are also expressed in *obscura* group testis (Figure S3), suggesting that if Ago2 paralogues have specialised to suppress TEs, they are doing so alongside the existing TE suppression mechanism. Second, testis-specificity could have evolved to suppress meiotic drive, which is prevalent (in the form of sex-ratio distortion) in the *obscura* group (Gershenson 1928; Sturtevant and Dobzhansky 1936; Wu and Beckenbach 1983; Jaenike 2001; Unckless et al. 2015), and which is suppressed by RNAi-based mechanisms in other species (Tao et al. 2007; Kotelnikov et al. 2009; Gell and Reenan 2013). A high level of meiotic drive in the *obscura* group could therefore impose selection for the evolution of novel suppression mechanisms, leading to the repeated specialization of Ago2 paralogues to the testis.

### Prospects for novel functions during the evolution of RNAi

The functional specialization that we observe for *obscura* group Ago2 paralogues raises the prospect of undiscovered derived functions following Argonaute expansions in other lineages. Ago2 has duplicated frequently across the arthropods, with expansions present in insects (*Drosophila willistoni* (Figure 1) & *Musca domestica*, Scott et al. 2014), crustaceans (*Penaeus monodon*, Leebonoi et al. 2015) and chelicerates (*Tetranychus urticae*, *Ixodes scapularis*, *Mesobuthus martensii* & *Parasteatoda tepidariorum*, Palmer and Jiggins 2015). The prevalence of testis-specificity in *obscura* group Ago2 paralogues raises the possibility that specialization to the germline may be more widespread following Argonaute duplication. The expression of Ago2 paralogues has previously been characterized in *P. monodon*, and shows that one paralogue has indeed specialised to the germline of both males and females, but not the testis alone (Leebonoi et al. 2015). Publicly available RNAseq data from the head, gonad and carcass of male and female *Musca domestica* (GSE67065, Meisel et al. 2015) suggests that neither *M. domestica* Ago2 paralogue has specialised to the testis (Figure S8). However, public data from the head, thorax and abdomen of male and female *D. willistoni* (GSE31723, Meisel et al. 2012) shows that one *D. willistoni* Ago2 paralogue (FBgn0212615) is expressed ubiquitously, while the other (FBgn0226485) is expressed only in the male abdomen (Figure S8), consistent with the evolution of testis-specificity after duplication. This raises the possibility that a testis-specific selection pressure may be driving the retention and specialization of Ago2 paralogues across the arthropods.

In conclusion, we have identified rapid and repeated evolution of testis-specificity after the duplication of Ago2 in the *obscura* group, associated with low genetic diversity and signatures of strong selection. Ago2 and other RNAi genes have undergone frequent expansions in different eukaryotic lineages (Mukherjee et al. 2013; Lewis et al. 2016), and have been shown to switch between ubiquitous and germline- or ovary-specific functions in isolated species. This study provides evidence for the evolution of a new testis-specific RNAi function, and suggests that positive selection may act on young paralogues to drive the rapid evolution of novel RNAi mechanisms across the eukaryotes.

## Acknowledgements

This work was supported by a Natural Environment Research Council Doctoral Training Grant (NERC DG NE/J500021/1 to SHL), the Academy of Finland (265971 to HS), a University of Edinburgh Chancellor’s Fellowship and a Wellcome Trust Research Career Development Fellowship (WT085064 to DJO), and a Wellcome Trust strategic award to the Centre for Immunity, Infection and Evolution (WT095831 to the CIIE). We thank Ben Longdon and Brian Charlesworth for providing us with strains of *D. obscura* and *D. pseudoobscura* respectively, and Francis Jiggins for providing us with DCV.

